# Stress-induced mitochondrial fragmentation in endothelial cells disrupts blood-retinal barrier integrity causing neurodegeneration

**DOI:** 10.1101/2024.12.21.629919

**Authors:** Jorge L. Cueva-Vargas, Nicolas Belforte, Isaac A. Vidal-Paredes, Florence Dotigny, Christine Vande Velde, Heberto Quintero, Adriana Di Polo

## Abstract

Increased vascular leakage and endothelial cell (EC) dysfunction are major features of neurodegenerative diseases. Here, we investigated the mechanisms leading to EC dysregulation and asked whether altered mitochondrial dynamics in ECs impinge on vascular barrier integrity and neurodegeneration. We show that ocular hypertension, a major risk factor to develop glaucoma, induced mitochondrial fragmentation in retinal capillary ECs accompanied by increased oxidative stress and ultrastructural defects. Analysis of EC mitochondrial components revealed overactivation of dynamin-related protein 1 (DRP1), a central regulator of mitochondrial fission, during glaucomatous damage. Pharmacological inhibition or EC-specific *in vivo* gene delivery of a dominant negative DRP1 mutant was sufficient to rescue mitochondrial volume, reduce vascular leakage, and increase expression of the tight junction claudin-5 (CLDN5). We further demonstrate that EC-targeted CLDN5 gene augmentation restored blood-retinal-barrier integrity, promoted neuronal survival, and improved light-evoked visual behaviors in glaucomatous mice. Our findings reveal that preserving mitochondrial homeostasis and EC function are valuable strategies to enhance neuroprotection and improve vision in glaucoma.

## INTRODUCTION

The blood-brain- and inner blood-retinal barriers (BBB, iBRB) regulate the transport of molecules in and out of the brain and retina, while preventing the entry of neurotoxic factors, immune cells, and pathogens^1, 2^. During development, endothelial cells (ECs) acquire key features to establish the BBB/iBRB, notably the expression of specialized tight junctions that restrict paracellular flow^3, 4^. Tight junction complexes in ECs contain integral membrane proteins including claudins, occludin, and junctional adhesion molecules, which mediate intercellular contacts and recruit cytoplasmic proteins^5^. ECs display a low rate of transcytosis that restricts vesicle-mediated transcellular movement of solutes^6^, while expressing transport systems that allow the exchange of nutrients and the elimination of toxins^7^. The integrity of the BBB and iBRB is critical to maintain an optimal environment for neuronal function and, as such, the loss of barrier integrity is a major component of prevalent brain and retinal neuropathologies^8^.

In the central nervous system, ECs have a relatively low mitochondrial volume accounting for 8-11% of their total cytoplasmic content^9^. The low mitochondrial volume found in ECs underlines the specialized role of mitochondria in signaling responses to environmental cues rather than solely energy production^10^. Indeed, ECs rely mostly on glycolysis instead of oxidative phosphorylation as their main energy source, with over 80% of ATP produced from glycolytic conversion of glucose to lactate^11^. Consistent with this, cumulative evidence support that EC mitochondria serve as signaling hubs that regulate many processes including gene expression, cellular growth, calcium homeostasis, response to hypoxic and oxidative stressors as well as hemodynamics^10, 12, 13^.

Mitochondria are endowed with a dynamic quality control system where they frequently fuse and divide to control their size, density, morphology, and distribution^14, 15, 16^. ECs maintain a healthy population of mitochondria by achieving a tightly regulated balance between fission and fusion^17^. The central regulator of mitochondrial fission is the GTPase dynamin-related protein 1 (DRP1), which is recruited from the cytosol to the outer mitochondrial membrane where it oligomerizes to form helical structures that encircle, constrict, and cleave the mitochondrion^18^. Fusion, on the other hand, is coordinated by mitofusins (MFN1, MFN2), located in the outer mitochondrial membrane, and optic atrophy protein 1 (OPA1) in the inner mitochondrial membrane^19^. Defective mitochondrial fusion and fission are linked to EC dysfunction and microvascular injury^20^. However, whether altered mitochondrial dynamics in ECs alters vascular permeability in neuropathologies is poorly understood. Here, we asked the following key questions: i) do altered mitochondrial dynamics contribute to loss of vascular integrity?, ii) if so, what are the mechanisms underlying mitochondria-related vessel leakage? and iii) what is the impact of mitochondrial alterations leading to barrier dysfunction on neurodegeneration?

To address this knowledge gap, we investigated *in vivo* mitochondrial alterations in the inner retinal microvasculature during ocular hypertension (OHT), a major risk factor for developing glaucoma, the leading cause of irreversible blindness worldwide^21^. Loss of vision in glaucoma is caused by the selective death of retinal ganglion cells (RGCs)^22^, a population of long projecting neurons that are highly vulnerable to microvascular dysfunction^23, 24^. Evidence from clinical and pre-clinical studies support that there is loss of vascular integrity during glaucomatous damage^23, 25, 26^, however, very little is known about the mechanisms underlying vessel leakage. Here, we demonstrate substantial mitochondrial fragmentation accompanied by oxidative stress and increased capillary leakage at the earliest stages of glaucomatous neurodegeneration. Our data indicate that overactivation of DRP1 during ocular pressure-induced damage increased mitochondrial fission in ECs. Furthermore, we show that blocking DRP1 function rescued mitochondrial volume, reduced vascular leakage, and increased claudin-5 (CLDN5) expression. Remarkably, adeno-associated virus (AAV)-mediated CLDN5 gene supplementation selectively in ECs improved iBRB integrity, promoted neuronal survival, and rescued light-evoked visual behaviors.

## RESULTS

### Retinal ECs undergo mitochondrial fragmentation during ocular pressure-dependent stress

To visualize mitochondria in the inner retinal vasculature we used transgenic mice carrying EGFP-labeled mitochondria driven by the mouse homeobox transcription factor Hb9 promoter (**Fig. 1A**), which is typically expressed by developing motoneurons^27^. Random integration of this transgene resulted in a founder with atypical EC-specific expression of EGFP-tagged mitochondria in the central nervous system, heart, spleen, thymus, lymph nodes, and skin^27^. We found that these mice, referred to as Endo-MitoEGFP, displayed robust EGFP labeling in EC mitochondria in the inner retina (**Fig. 1B**), across all vascular plexuses, which was confirmed by co-labeling with Cytochrome c (Cytc) as well as the vascular marker CD31 and lectin (**Fig. 1C-J**). To mimic OHT-dependent glaucomatous stress, Endo-MitoEGFP mice received an intracameral injection of magnetic microbeads, which were attracted to the iridocorneal angle with a magnet to block aqueous humor outflow (**Supp. Fig. 1A**). This procedure leads to gradual increase in intraocular pressure and progressive RGC death, key landmarks of pathological changes in glaucoma^28^ (**Supp. Fig. 1B-D**). Sham controls received an intracameral injection of vehicle without microbeads. Mitochondrial volume was evaluated at three weeks after OHT induction (OHT-3wk) because intraocular pressure increase is stable at this timepoint and RGC loss is modest^28, 29^. Three-dimensional (3D) reconstruction of mitochondrial complexes in capillary ECs showed marked reduction of mitochondrial volume in eyes with high intraocular pressure (**Fig. 1K-Q**). We classified mitochondrial complexes in retinal ECs into three populations based on the quartiles of the volume distribution in sham retinas: large (>1 µm^3^), medium (0.1-1 µm^3^), and small (<0.1 µm^3^) (**Supp. Fig. 1E**). We found a significant reduction in the number of large mitochondria in eyes subjected to OHT relative to sham-operated controls (sham: 31%, OHT: 11.5%), whereas small mitochondria increased proportionally (sham: 11%, OHT: 33%), and medium-sized complexes remained unaffected (**Fig. 1R-T**). To rule out that these changes resulted from capillary dropout, we quantified microvascular density across all vascular plexuses and found that capillary density was the same in sham and OHT-exposed retinas (**Supp. Fig. 2**).

**Figure 1.**
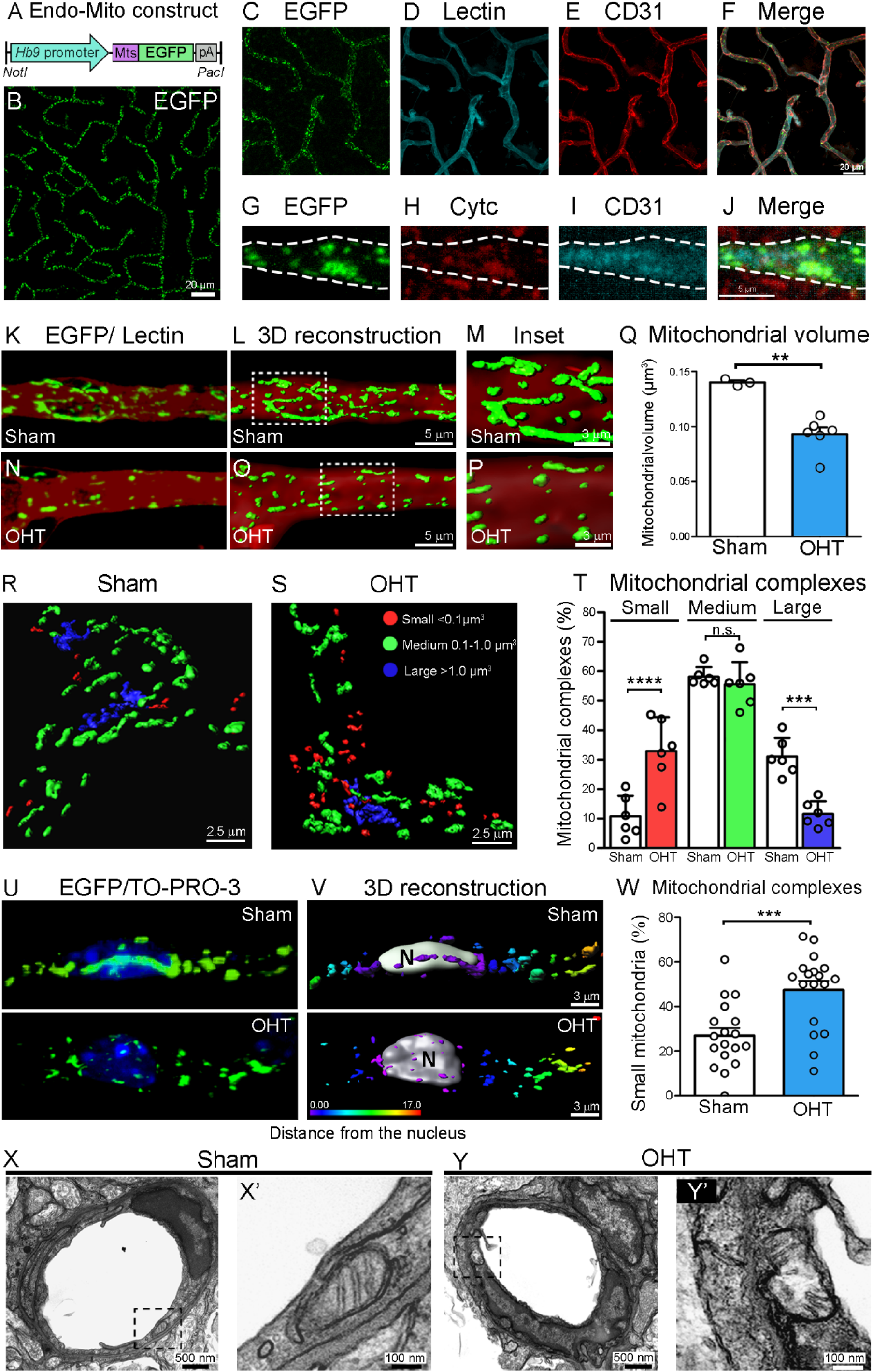
Retinal ECs undergo mitochondrial fragmentation during ocular pressure-dependent stress. (A) Schematic of the construct used to generate Endo-MitoEGFP mice including the mitochondrial targeting sequence (Mts) of Cytochrome c (Cytc) oxidase subunit VIII. (B-F) Whole-mount retinal preparations demonstrate the presence of GFP-tagged mitochondria in retinal capillaries, visualized with lectin and the endothelial cell (EC) marker CD31. (G-J) Representative retinal cross-section shows co-localization of EGFP-labeled mitochondria with Cytc in ECs identified with CD31 (dotted line shows capillary outline). (K-P) 3D reconstruction of mitochondrial complexes within retinal capillaries shows decreased mitochondrial volume at 3 weeks of ocular hypertension (OHT) induction. Insets in (L) and (O) show magnified areas in (M) and (P), respectively. (Q) Quantitative analysis confirms that OHT causes loss of mitochondrial volume (sham: 100%, OHT: 66%, two-tailed Student’s t-test, N= 4-6 mice/group, **p< 0.01). (R-S) 3D reconstruction of mitochondrial complexes reveals three populations based on volume distribution: large (>1 µm^3^, blue), medium (0.1-1 µm^3^, green), and small (<0.1 µm^3^, red). (T) The number of large mitochondria in OHT eyes decrease whereas small mitochondria increase, and medium-sized complexes remain unaffected (N=6 mice/group, two-way ANOVA, Sidak’s multiple comparison *post-hoc* test, ***p=0.0002, ****p<0.0001). (U, V) Mitochondrial fragmentation in the perinuclear area of ECs subjected to OHT. Color coding denotes distance from nucleus (N=nucleus). (W) Quantitative analysis shows a higher number of small perinuclear mitochondrial complexes in glaucomatous retinas (OHT: 47.5%, sham: 27%, N=6 mice/group, two-tailed Student’s t-test, ***p< 0.001). (X, Y) TEM confirms ultrastructural alterations in EC mitochondria caused by OHT. Data are presented as mean values ± S.E.M.

Mitochondria are known to communicate with the nucleus to coordinate changes in gene expression, metabolism, and stress-related adaptation^30^. As such, mitochondria exist in proximity to the nucleus and can establish direct mitochondrial-nuclear associations as a result of stress from mitochondrial reactive oxygen species (mtROS) leading to transcriptional changes^31, 32^. Analysis of perinuclear mitochondrial complexes revealed an increase in mitochondrial fragmentation in glaucomatous retinas relative to sham controls (**Fig. 1U-W**). Our data show a ∼2-fold increase in the number of small mitochondrial complexes surrounding EC nuclei in retinas subjected to OHT (48%) compared to controls (26%) (**Fig. 1W**). To investigate whether changes in mitochondrial volume were associated with ultrastructural alterations in EC mitochondria, we carried out transmission electron microscopy (TEM). We found abnormalities in mitochondria morphology during OHT including disrupted membranes and cristae (**Fig. 1X, 1Y**). These results demonstrate that OHT-induced stress leads to mitochondrial fragmentation and ultrastructural alterations in retinal capillary ECs.

### ECs with fragmented mitochondria display overactive DRP1, oxidative stress, and increased vascular leakage

To assess the potential contribution of disrupted EC mitochondrial dynamics to glaucomatous damage, we first examined the levels of DRP1, MFN1, MFN2, and OPA-1 in retinal ECs. Phosphorylation of DRP1 plays a crucial role in DRP1 activity regulation. Specifically, DRP1 phosphorylation at Serine 616 (pDRP1^Ser616^) promotes the translocation of DRP1 from the cytosol to the mitochondrial outer membrane where it triggers mitochondrial fission^33^. Western blot analysis of enriched retinal EC lysates from eyes subjected to three weeks of OHT revealed an increase in pDRP1^Ser616^ and total DRP1 relative to sham-operated controls (**Fig. 2A**). The levels of MFN1, MFN2, and OPA-1 remained unchanged (**Fig. 2B-D**). Mitochondrial fragmentation induces excess mtROS that can result in EC dysfunction^34^. To examine this, we performed an intravenous injection of MitoSOX-Red in Endo-MitoEGFP mice to measure the levels of mitochondrial superoxide in retinal ECs. Our data show a 2.3-fold increase in superoxide levels in EC mitochondria in glaucomatous retinas compared to sham controls (**Fig. 2E-K**).

**Figure 2.**
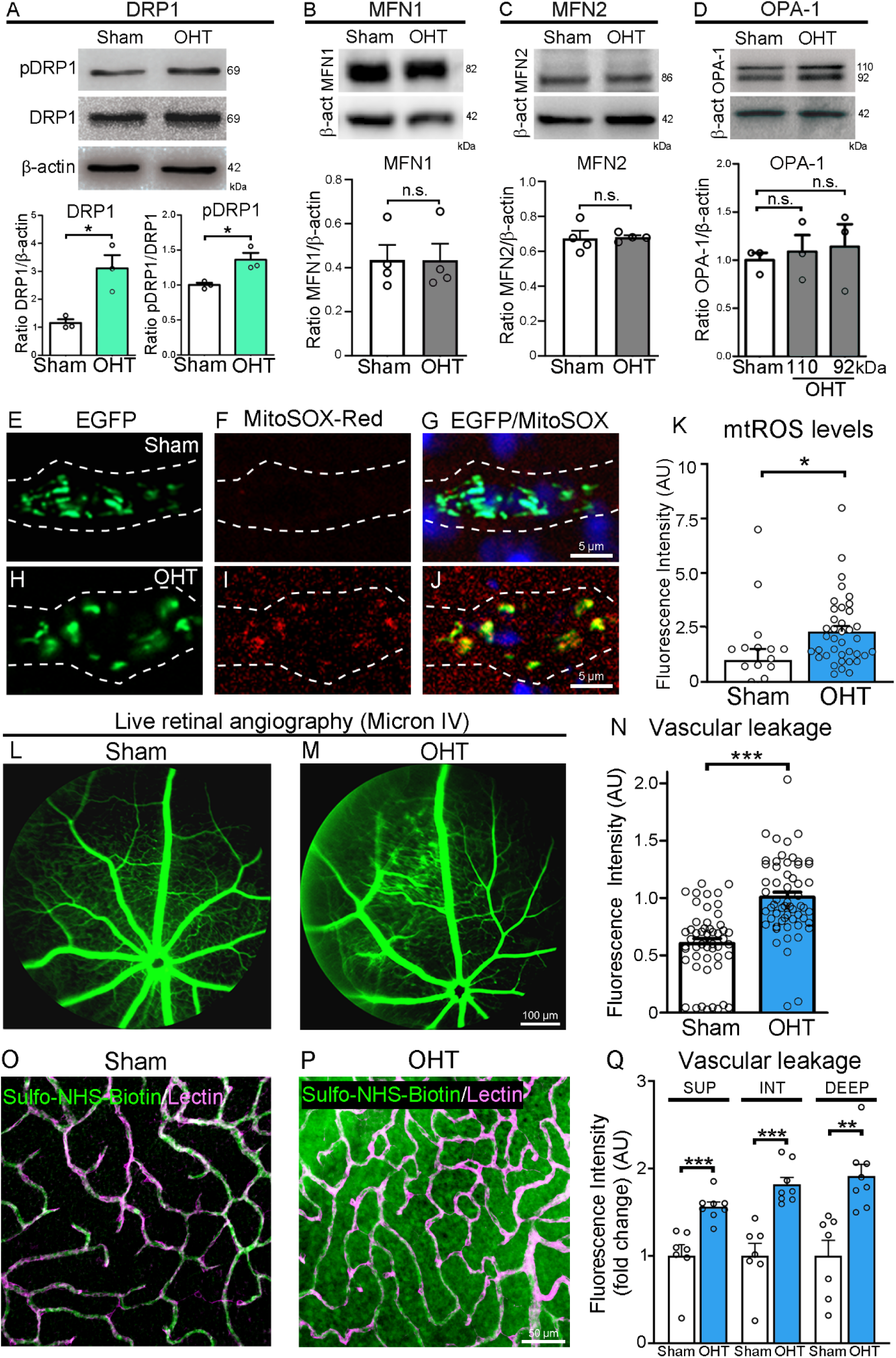
ECs with fragmented mitochondria display overactive DRP1, oxidative stress, and increased vascular leakage. (A-D) Western blot analyses of enriched retinal EC preparations show an increase in phosphorylated DRP1 at Serine 616 (pDRP1, active form), whereas MFN1, MFN2, and OPA-1 remained unchanged (N=3 mice/group, two-tailed Student’s t-test, \**p*<0.05, n.s.: not significant). Total DRP1 and/or b-actin were used as internal controls. (E-J) Whole-mount retinal preparations labeled with MitoSOX-Red show increased mitochondrial superoxide in ECs subjected to OHT. (K) Quantitative analysis confirm a 2.3-fold increase in superoxide levels in EC mitochondria from glaucomatous retinas compared to sham controls (N=4 mice/group, two-tailed Student’s t-test; *p<0.05). (L-N) *In vivo* fluorescein angiography images show tracer leakage in eyes subjected to glaucomatous damage (N=4 mice/group, two-tailed Student’s t-test, ***p<0.001). (O, P) Representative flat-mounted retinas show Sulfo-NHS-LC-Biotin leakage in eyes with OHT compared to sham controls. (Q) Quantification of Sulfo-NHS-LC-Biotin in the parenchyma confirms increased capillary permeability in the superficial (SUP), intermediate (INT), and deep (DEEP) vascular plexuses (N=8 mice/group, **p<0.005, ***p<0.001). Data are presented as mean values ± S.E.M.

Next, we examined whether mitochondrial fragmentation and increased mtROS production were associated with loss of iBRB integrity using two complementary leakage assays: i) live retinal angiography imaging with fluorescein (molecular weight: 332.32 Da), and ii) stereological analysis of the retinal capillary network using Sulfo-NHS-LC-Biotin (molecular weight: 443.43 Da), a tracer that binds to amine-containing molecules including cell surface proteins^35, 36^. After intravenous injection, fluorescein was confined to vessels in sham-operated control mice, whereas it leaked out of capillaries in mice subjected to glaucomatous damage (OHT-3wk) (**Fig. 2L-N**). Leaky hotspots of fluorescein were apparent throughout OHT-stressed retinas **(Fig. 2M**). To validate these findings, we examined barrier integrity after retro-orbital injection of Sulfo-NHS-LC-Biotin. We found that Sulfo-NHS-LC-Biotin remained inside retinal capillaries in sham control mice but leaked to the parenchyma in mice with glaucoma (**Fig. 2O, 2P**). Diffuse leakage was detected in all retinal quadrants and vascular plexuses (superficial, intermediate, and deep) (**Fig. 2Q**). Collectively, these data demonstrate DRP1 overactivation together with increased oxidative stress in ECs and loss of vascular integrity at the early stages of glaucomatous damage.

### DRP1 pharmacological inhibition replenishes mitochondria, reduces mtROS, restores iBRB function, and promotes neuroprotection

DRP1-dependent mitochondrial fission has been shown to play a critical role in cardiovascular diseases and EC dysfunction^37^. To investigate the functional contribution of DRP1 to mitochondrial alterations and vascular leakage in glaucoma, we first used the pharmacological inhibitor Mitochondrial Division Inhibitor-1 (Mdivi-1), a small molecule that crosses the BBB/BRB barriers, to block DRP1 function^38, 39^. Endo-MitoEGFP mice received daily intraperitoneal injections of Mdivi-1 starting at one week after OHT induction and mitochondrial volume and mtROS were examined two weeks later (OHT-3wk) (**Fig. 3A**). Midivi-1 prevented EC mitochondrial fragmentation and replenished organelle volume in glaucomatous retinas relative to vehicle-treated controls (**Fig. 3B, 3C**). We found a substantial decrease in the number of fragmented small mitochondrial complexes accompanied by an increase in large complexes in retinas from mice treated with Mdivi-1 (**Fig. 3D**). The number of medium-sized mitochondrial complexes did not change (**Supp. Fig. 3A**). Moreover, quantitative analysis of mtROS with MitoSOX-Red demonstrated a marked reduction in oxidative stress following Mdivi-1 administration compared to vehicle-treated controls (**Fig. 3E-K**).

**Figure 3.**
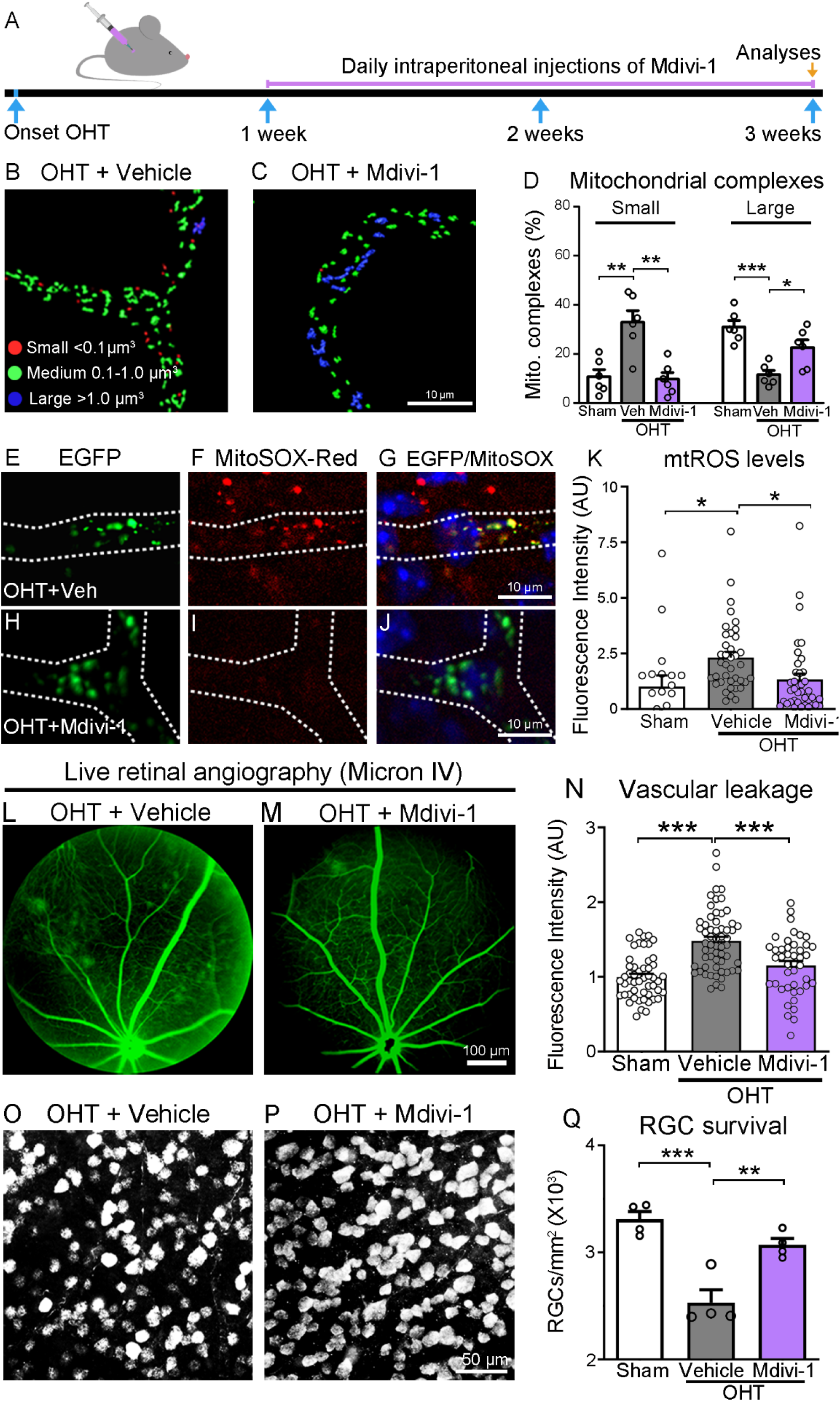
DRP1 pharmacological inhibition replenishes mitochondria, reduces mtROS, restores iBRB function, and promotes neuroprotection. (A) Timeline of Mdivi-1 treatment and analyses. (B-D) 3D reconstruction of mitochondrial complexes show that Mdivi-1 prevented EC mitochondrial fragmentation and replenished organelle volume in retinas subjected to OHT relative to vehicle-treated controls (N=6 mice/group, ANOVA with Tukey’s multiple comparison *post-hoc* test, \**p*<0.05, \*\**p*<0.005, ***p<0.001). (E-J) Representative images from flat-mounted retinas show decreased MitoSOX-Red labeling in ECs after Midivi-1 treatment. Dotted lines show vessel outline. (K) Quantitative analysis confirms reduced oxidative stress in EC mitochondria following Mdivi-1 treatment relative to vehicle-treated controls (N=4 mice/group, ANOVA with Tukey’s multiple comparison *post-hoc* test; *p<0.05). (L-N) *In vivo* fluorescein angiography images show that Midivi-1 reduces tracer leakage in glaucomatous eyes compared to controls. (N) Quantification of fluorescein intensity confirms that Mdivi-1 preserves the iBRB integrity during OHT-induced damage relative to vehicle-treated eyes (N=4 mice/group, ANOVA with Tukey’s multiple comparison *post-hoc* test, ***p<0.001). (O, P) Representative flat-mounted retinas labeled with the RGC-specific marker RBPMS show higher neuronal density following Mdivi-1 treatment. (Q) Quantification of RGC density confirms that Mdivi-1 promotes significant neuroprotection (N=8 mice/group, ANOVA with Tukey’s multiple comparison *post-hoc* test; \*\**p*<0.005, \*\*\**p*<0.001). Data are presented as mean values ± S.E.M. The cartoon in this figure was generated with BioRender (https://biorender.com).

To investigate whether DRP1 inhibition had an impact on barrier integrity, we performed fluorescein angiography in eyes subjected to OHT and Mdivi-1 or vehicle treatment. Our data show a significant decrease in tracer leakage in Mdivi-1-treated eyes relative to controls (**Fig. 3L-N**). Next, we asked whether improved EC mitochondrial dynamics with Mdivi-1 impacted RGC survival during OHT. For this purpose, Mdivi-1 was administered using the same regimen shown in **Figure 3A** and RGC density was quantified at 3 weeks after OHT induction, a time when there is sufficient RGC loss in our glaucoma model (∼20-25%) to allow the assessment of neuroprotection (**Supp. Fig. 1**)^28, 29, 40, 41^. RGCs were visualized with the cell-specific marker RBPMS (RNA-Binding Protein with Multiple Splicing)^42^. Mdivi-1 promoted RGC survival and supported neuronal density to levels similar to those in uninjured sham-operated control (**Fig. 3O-Q**). Mdivi-1 treatment did not reduce mean intraocular pressure over the duration of the study (Midivi-1= 20.6±1.1 mm Hg; vehicle= 20.1±0.9 mm Hg, N=7mice/group, two-tailed Student’s t-test, p=0.92), thus its effect cannot be attributed to OHT lowering. Together, these findings demonstrate that pharmacological inhibition of DRP1 replenishes EC mitochondria, attenuates oxidative stress, rescues barrier function, and promotes neuronal survival.

### EC-specific gene transfer of dominant negative DRP1 restores barrier integrity, improves neuronal survival, and increases expression of CLDN5

To rule out pan-retinal and off-target effects of pharmacological DRP1 inhibition, we sought to decrease DRP1 function selectively in ECs by reducing the GTPase activity of DRP1 using a dominant-negative mutant (DRP1^K38A^)^43^. For this purpose, we used a serotype 9 adeno-associated virus encoding DRP1^K38A^ (AAV.DRP1^K38A^) under control of the Ple32 minipromoter, a small regulatory sequence based on the *CLDN5* gene promoter that drives selective gene expression in retinal ECs^44^. Endo-MitoEGFP mice received a single tail vein injection of AAV.DRP1^K38A^ three weeks prior to OHT induction and mitochondrial fragmentation, mtROS, and fluorescein leakage were analyzed over the course of the following three weeks (**Fig. 4A**). Control mice received the same AAV construct but lacking DRP1^K38A^ (AAV.Ctl). There was no statistically significant difference in elevated mean intraocular pressure in glaucomatous mice treated with AAV.DRP1^K38A^ or AAV.Ctl (AAV.DRP1^K38A^=22.4±1.2 mm Hg; AAV.Ctl = 20.1±0.9 mm Hg, N=7 mice/group, two-tailed Student’s t-test, p=0.23). Analysis of mitochondrial volume revealed that AAV-mediated DRP1^K38A^ reduced the number of small mitochondrial complexes while increasing the frequency of large complexes (**Fig. 4B-D**) indicating that attenuation of DRP1 function reduced mitochondrial fission in ECs. These changes in mitochondrial volume were accompanied by reduced mtROS levels, detected using MitoSOX-Red, in eyes treated with AAV.DRP1^K38A^ (**Fig. 4E-K**). Importantly, our data show that selective expression of dominant negative DRP1 in ECs by AAV.DRP1^K38A^ markedly reduced fluorescein leakage throughout the retina (**Fig. 4L-N**) and promoted robust RGC survival in glaucomatous eyes relative to AAV.Ctl (**Fig. 4O-Q**).

**Figure 4.**
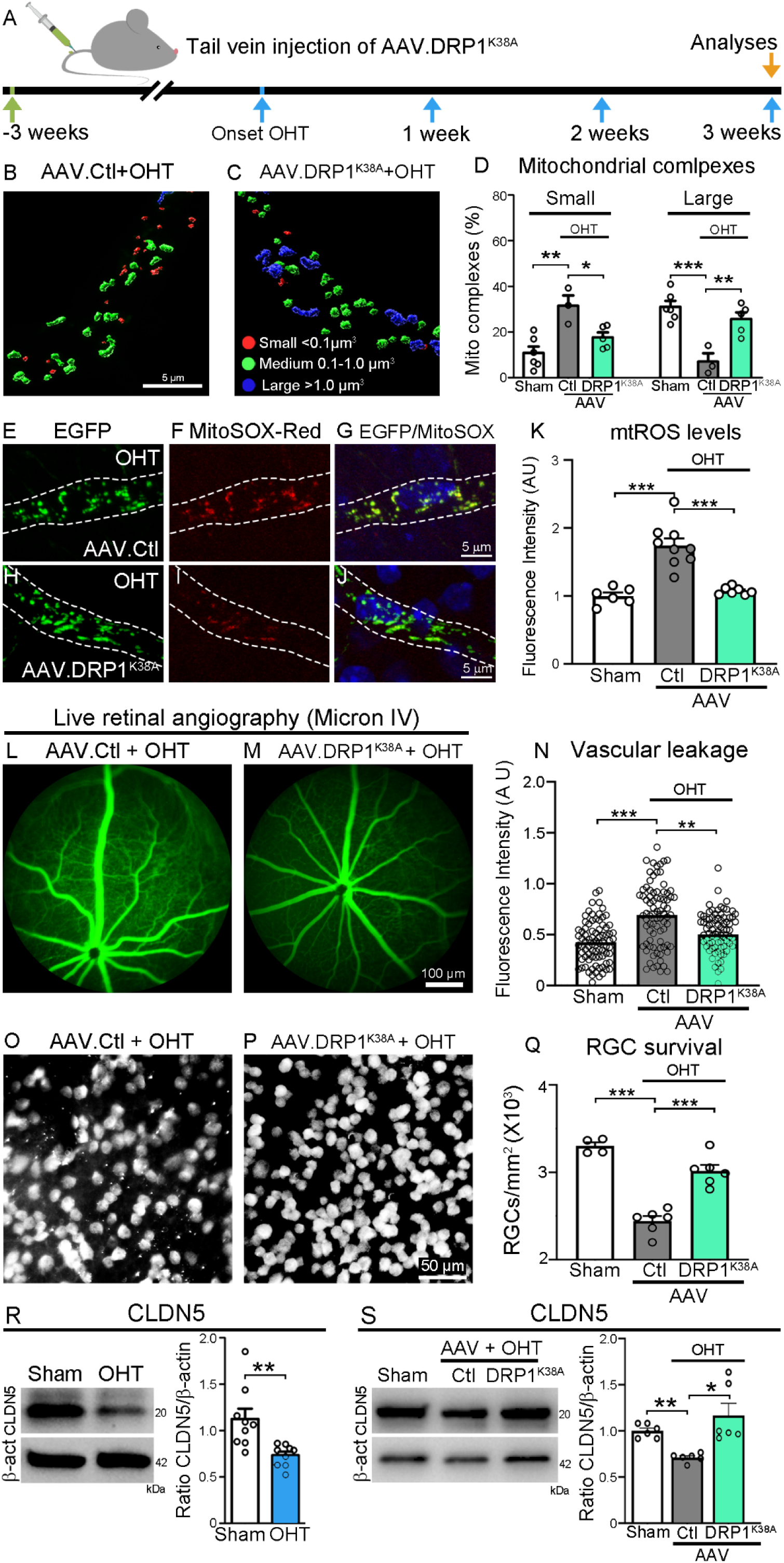
EC-specific gene transfer of dominant negative DRP1 restores barrier integrity, improves neuronal survival, and increases expression of CLDN5. (A) Timeline for administration of AAV.DRP1^K38A^ and subsequent analyses. (B-D) 3D reconstruction of mitochondrial complexes show that AAV.DRP1^K38A^ restores mitochondrial volume in ECs (N=3-6 mice/group, ANOVA with Tukey’s multiple comparison *post-hoc* test, *p<0.05, **p<0.01, ***p<0.001). (E-K) Inhibition of DRP1 with AAV.DRP1^K38A^ decreases oxidative stress in EC mitochondria in glaucomatous retinas assessed with MitoSOX-Red (N=4 mice/group, ANOVA with Tukey’s multiple comparison *post-hoc* test, ***p<0.0001). (L-N) *In vivo* fluorescein angiography shows that AAV.DRP1^K38A^ preserves the iBRB integrity in glaucomatous eyes compared to mice treated with AAV.Ctl (N=4 mice/group, ANOVA with Tukey’s multiple comparison *post-hoc* test, **p<0.01, ***p<0.001). (O-Q) Restoration of vascular barrier integrity with AAV.DRP1^K38A^ resulted in robust neuroprotection based on quantification of RBPMS-labeled RGC density (N=8 mice/group, with Tukey’s multiple comparison *post-hoc* test; ***p<0.0001). (R) Western blot analysis of fresh retinal lysates show that CLDN5 expression is downregulated in glaucomatous retinas compared to sham-operated controls (N=9 mice/group, Student’s t-test, \*\**p*<0.01). (S) Western blot analysis shows that AAV.DRP1^K38A^ treatment in glaucomatous eyes restores CLDN5 to pre-injury (sham) levels (N=6-9 mice/group, ANOVA with Tukey’s multiple comparison *post-hoc* test, *p<0.05, **p<0.01). Data are presented as mean values ± S.E.M. The cartoon in this figure were generated with BioRender (https://biorender.com).

Low molecular weight tracers like fluorescein and biotin are known to diffuse into the parenchyma via paracellular pathways upon EC tight junction disruption^35, 36^. Based on this, we investigated OHT-induced changes in the tight junction proteins CLDN5, occludin, and zona occludens-1 (ZO-1). We found a marked reduction of CLDN5 in glaucomatous retinas relative to sham-operated controls (**Fig. 4R**), whereas occludin and ZO-1 remained unchanged (**Supp. Fig. 4A. 4B**). In addition, the levels of the adherens junctions vascular endothelial cadherin (VE-cadherin), notably its phosphorylated active forms (pCadh^Thr731^ and pCadh^Thr685^), and b-catenin did not change with OHT induction (**Supp. Fig. 4C-E**). CLDN5 is selectively expressed by ECs^45^ and is highly enriched in the retinal microvasculature^46, 47^. EC mitochondria, through changes in mtROS and physical proximity to the nucleus, can regulate gene transcription and protein expression^31, 32^. Therefore, we asked whether reduced DRP1-dependent mitochondrial fission altered CLDN5 levels. Western blot analyses of retinal extracts demonstrated that glaucomatous retinas treated with AAV.Ctl displayed marked reduction of CLDN5, whereas inhibition of DRP1 with AAV.DRP1^K38A^ restored CLDN5 to sham-like levels (**Fig. 4S**). Taken together, these data demonstrate that EC-targeted inhibition of DRP1 function improves barrier integrity and neuronal survival in glaucoma, a response that is associated with increased CLDN5 expression.

### CLDN5 gene therapy preserves BRB integrity, enhances neuronal survival, and rescues visual function in glaucoma

Based on the effect of DRP1 inhibition on EC function and increased CLDN5 expression, we then asked whether *CLDN5* gene supplementation had an impact on barrier integrity and neurodegeneration during glaucomatous damage. To this end, we generated a serotype 9 AAV encoding CLDN5 under control of the Ple32 minipromoter (AAV.CLDN5) to selectively upregulate CLDN5 in retinal ECs (**Fig. 5A**). Analysis of retinas from Endo-MitoEGFP mice that received a tail vein injection of AAV.CLDN5, using the regimen shown in **Figure 4A**, confirmed selective expression of CLDN5 in ECs (**Fig. 5B-D**). No other retinal cell types displayed CLDN5 expression after tail vein injection of AAV.CLDN5 (**Supp. Fig. 5A**). Next, we examined the effect of CLDN5 gene supplementation on vascular integrity using fluorescein angiography, and found that AAV.CLDN5 substantially reduced tracer leakage in eyes subjected to OHT relative to controls (**Fig. 5E-G**). To further validate these findings, we performed retro-orbital injections of Sulfo-NHS-LC-Biotin and analyzed tracer leakage in all vascular plexuses using *ex vivo* retinal preparations. Glaucomatous eyes exposed to control AAV.Ctl showed an increase of Sulfo-NHS-LC-Biotin fluorescence in the parenchyma, whereas eyes treated with AAV.CLDN5 showed reduced vascular permeability (**Fig. 5H, I**). Quantitative analysis confirmed that AAV.CLDN5 reduced capillary leakage in glaucoma, across vascular plexuses, to levels like those found in sham-operated mice (**Fig. 5J**).

**Figure 5.**
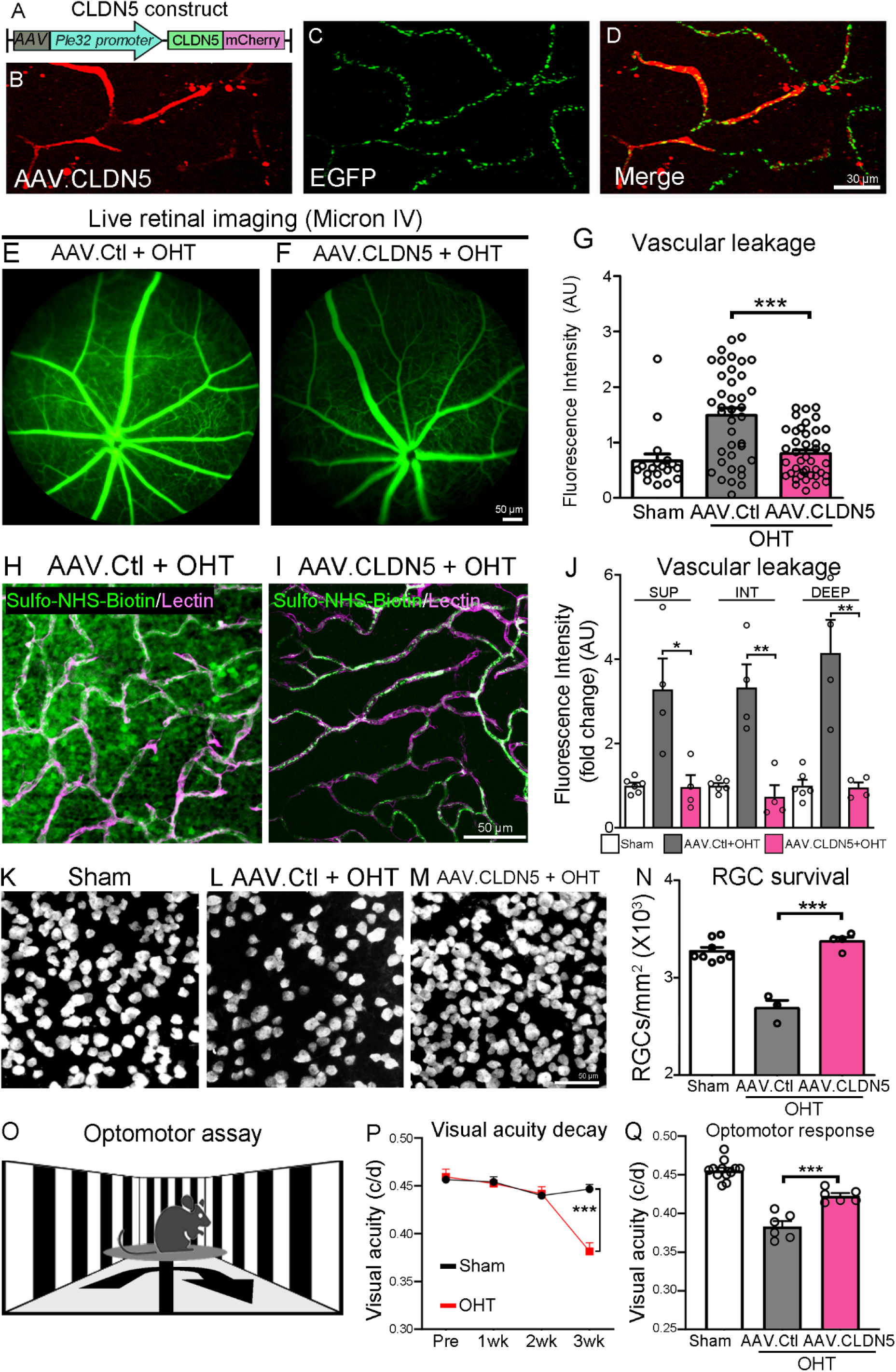
CLDN5 gene therapy preserves BRB integrity, enhances neuronal survival, and rescues visual function in glaucoma. (A) Construct used to generate AAV serotype 9 carrying the *CLDN5* gene with an mCherry tag under control of the Ple32 (*CLDN5*) promoter (AAV.CDLN5). (B-D) AAV-mediated CLDN5-mCherry expression in retinal ECs (B) co-labeled with EGFP-tagged mitochondria (C) in Endo-Mito mice. (E-G) *In vivo* fluorescein angiography imaging shows that AAV-mediated CLDN5 supplementation reduces iBRB permeability in glaucomatous retinas relative to controls (AAV.Ctl) (N=4 mice/group, ANOVA with Tukey’s multiple comparison *post-hoc* test, ***p<0.001). (H, I) Representative confocal micrographs show reduced Sulfo-NHS-LC-Biotin vascular leakage during OHT-induced damage after AAV.CLDN5 treatment. (J) Quantitative analysis confirms that retinal barrier integrity is markedly improved in all vascular plexuses (SUP, INT, DEEP) in AAV.CLDN5-treated glaucomatous eyes relative to AAV.Ctl-treated controls (N=5 mice/group, ANOVA with Tukey’s multiple comparison *post-hoc* test, *p<0.05, \*\**p*<0.001). (K-M) Representative images of RBPMS-labeled flat-mounted retinas from sham (K) and 3 weeks of OHT treated with AAV.Ctl (L) or AAV.CLDN5 (M). (N) Quantification of RGC density shows that AAV.CLDN5 promotes neuronal survival at 3 weeks of OHT induction relative to control eyes (N=4-8 mice/group, ANOVA with Tukey’s multiple comparison *post-hoc* test, ***p<0.001) (O) Optomotor reflex assay setup. (P) Longitudinal evaluation of the optomotor reflex before (Pre) and after magnetic microbead injection demonstrates progressive visual acuity decay resulting in significant vision loss by 3 weeks of exposure to OHT (N=6-7 mice/group, two-way ANOVA with Sidak’s multiple comparison *post hoc* test, ***p<0.001) (c/d=cycles/degree). (Q) Quantitative analysis of the optomotor responses at 3 weeks post-OHT induction shows a significant improvement in visual acuity in mice treated with AAV.CLDN5 relative to AAV.Ctl (N=6-12 mice/group, ANOVA with Tukey’s multiple comparison *post-hoc* test, ***p<0.001). Data are presented as mean values ± S.E.M. The cartoon in this figure were generated with BioRender (https://biorender.com).

Next, we evaluated the effect of CLDN5 gene therapy on neuronal survival by quantifying RGC density after AAV.CLDN5 or AAV.Ctl administration to assess neuroprotection at three weeks of OHT. Robust RGC survival was observed after AAV.CLDN5 treatment, which supported neuronal density at levels found in sham-operated control eyes (non-injured, 100% survival), while RGC loss was observed in control retinas treated with AAV.Ctl (**Fig. 5K-N**). AAV.CLDN5 administration did not reduce OHT (AAV.CLDN5=20.3±1.2 mm Hg; AAV.Ctl= 20.2±1.2 mm Hg, N=7 mice/group, two-tailed Student’s t-test, p=0.74), thus neuroprotection could not be attributed to intraocular pressure lowering. To assess retina-brain connectivity we measured the optomotor reflex response, a visual behavior characterized by direction-selective head movements triggered by RGC inputs^48^. Mice from all groups were subjected to longitudinal (weekly) evaluation of the stereotyped behavior evoked by grating light stimuli and visual acuity was calculated (**Fig. 5O, 5P**). Visual acuity decreased in a time-dependent manner in eyes with OHT and became statistically lower than sham controls starting at three weeks after glaucoma induction (**Fig. 5P**), hence we evaluated the effect of CLDN5 gene transfer at this timepoint. Our data show that AAV.CLDN5 improved visual acuity in glaucomatous mice compared to AAV.Ctl-treated controls (**Fig. 5Q**). Together, these results reveal major beneficial effects of CLDN5 gene augmentation including preservation of iBRB integrity, enhancement of neuronal survival, and improvement of visual responses.

## DISCUSSION

The breakdown of vascular barriers in retinal and brain pathologies results in leakage of harmful components from the blood leading to inflammation, cellular infiltration, aberrant clearance, and neuronal deficits^49^. In glaucoma, clinical studies showed fluorescein filling defects and increased vascular permeability in patients with high intraocular pressure relative to normotensive individuals^23^. Within ECs, mitochondria integrate environmental cues including oxygen, nutrients, hemodynamics, and stress signals to modify cellular morphology and function^10^. How are EC mitochondria altered during glaucomatous damage, and do they impact barrier integrity and neurodegeneration? Our study reports several novel findings including: i) OHT-induced stress induces mitochondrial fission in retinal ECs leading to vascular leakage; ii) these alterations are rescued by blocking DRP1 function, which replenishes mitochondrial volume, reduces oxidative stress, decreases barrier permeability, and enhances CLDN5 expression; and iii) EC-targeted CLDN5 gene supplementation is sufficient to restore vascular integrity, promote neuronal survival, and improve vision.

To investigate structural changes in EC mitochondria during OHT-dependent stress, we used Endo-MitoEGFP mice, a founder line with a novel vascular-specific expression pattern created by random integration of a transgene encoding mitochondrially-targeted EGFP driven by the *Hb9* promoter^27^. Mitochondria isolated from brain, spinal cord, and spleen of Endo-MitoEGFP mice demonstrated EGFP localization to fractions enriched for mitochondria^27^. Here, we found EC-specific mitochondrial EGFP expression in the Endo-MitoEGFP retina and demonstrate the utility of this model to evaluate mitochondria structure and function in the visual system. We describe three distinct populations of mitochondrial complexes in retinal ECs: large (>1 µm^3^), medium (0.1-1 µm^3^), and small (<0.1 µm^3^) that are differentially affected by high intraocular pressure. In eyes subjected to OHT, the number of small mitochondrial complexes increased, notably in the perinuclear region, whereas large mitochondria decreased proportionately. Mitochondrial volume homeostasis not only plays a crucial housekeeping role in maintaining the structural integrity of this organelle but also affects many cellular functions. For example, an increase in mitochondrial matrix volume is sufficient to enhance fatty acid oxidation, respiration and ATP production^50^, whereas mitochondrial damage and reduced volume are associated with ROS production^51^. Consistent with this, we found increased mtROS in fragmented EC mitochondria from glaucomatous eyes than controls. These changes, together with DRP1 overactivation and their reversibility with strategies that blocked DRP1 function, strongly suggested heightened mitochondrial fission in ECs during glaucomatous stress.

The balance of fusion and fission is essential to maintain the quality and function of mitochondria and ensure normal cellular processes. In physiological conditions, mitochondrial fission helps divide damaged mitochondria into smaller organelles that can be targeted for elimination by mitophagy^52^. However, excessive mitochondrial fission is prevalent in stressed cells and is associated with mitochondrial dysfunction in a number of pathologies^53^. Although many studies have focused on altered mitochondrial dynamics in neurons and other cells including cardiomyocytes^54, 55^, far fewer reports exist on the impact of mitochondrial fission on ECs^10^. For example, mitochondrial fragmentation in RGCs has been linked to neuronal damage^56, 57, 58, 59^, but a role for mitochondrial fission in ECs during glaucomatous damage has not been reported. Our data show that increased DRP1 phosphorylation at serine 616 (pDRP1^Ser616^), which results in overactive DRP1, promotes mitochondrial fission and mtROS production in retinal ECs during OHT stress. In human umbilical vein ECs, sulfenylation of DRP1 induced mitochondrial fragmentation and mtROS elevation driving EC senescence and angiogenesis^34^.

We demonstrate that OHT-induced mitochondrial abnormalities led to EC dysfunction manifested primarily by the loss of barrier integrity across vascular plexuses. Indeed, targeted expression of dominant negative DRP1 exclusively in ECs or pharmacological DRP1 inhibition rescued mitochondrial defects and reduced iBRB permeability. Critically, EC-specific DRP1 inhibition promoted RGC survival indicating that EC dysfunction leading to vascular leakage plays a detrimental role during glaucomatous neurodegeneration. Our data support a model in which eye pressure-dependent stress alters ECs leading to DRP1 overactivation, mitochondrial fission and EC dysfunction. EC mitochondria are known to respond to oxygen and nutrient levels as well as hemodynamics^10^. Recent reports demonstrated compromised blood flow and neurovascular coupling at the early stages of OHT-induced damage^24^, thus it is possible that reduced oxygenation, low metabolites, and/or changes in laminar shear stress^60^ activate signaling pathways that recruit DRP1 leading to mitochondrial fission and EC dysfunction.

Our analysis of mitochondrial volume revealed substantial mitochondrial fission in the perinuclear region of ECs from retinas exposed to OHT. Overexpression of hFis1, a protein that promotes mitochondrial fission by recruiting DRP1, induces rapid fragmentation of mitochondria as well as clustering around the cell nucleus^61^. Perinuclear mitochondria is involved in ROS balance, which contributes to the transcriptional regulation of hypoxia-sensitive genes^61, 62^. Furthermore, an increase in intracellular ROS caused by dysfunctional mitochondria can serve as a signal to attenuate protein synthesis^63^. Based on this, we asked whether DRP1-dependent mitochondrial fission and mtROS production altered the levels of key components of the iBRB. We found that CLDN5 is markedly reduced in glaucomatous eyes, whereas other tight and adherens junctions remained unchanged. Reduced CLDN5 levels cause profound impairment of barrier integrity. For example, *CLDN5^−/−^* mice die at birth^64^ and *CLDN5* knockdown mice display severe disruption of the BBB with acute neuroinflammation and cell extravasation^65^. Recent data linked low CLDN5 levels with compromised BBB in neurodegenerative and neuropsychiatric disorders^66^. DNA methylation of the *CLDN5* gene has been observed in brain regions of Alzheimer’s disease patients and was associated with cognitive decline^67^. CLDN-5 expression is reduced in the hippocampus of patients with schizophrenia^68^, and mutations in CLDN-5 correlate with the development of epilepsy^69^. Importantly, our study shows that blocking DRP1 function rescued CLDN5 levels in retinal ECs, suggesting a critical role of CLDN5 in the loss of vascular integrity in glaucoma.

We report a strategy aimed at targeting the vasculature to enhance CLDN5 levels using AAV serotype 9 and a novel minipromoter (Ple32)^44^ to drive EC-specific *CLDN5* expression (AAV.CLDN5). Our data show that EC-targeted CLDN5 gene supplementation was sufficient to stabilize the iBRB and block vascular leakage, promote RGC survival, and improve visual acuity. These results not only confirm that CLDN5 is essential to maintain iBRB integrity, but also provide direct evidence that vascular leakage has a detrimental effect on neuronal survival and function. Remarkably, AAV.CLDN5 treatment promoted complete RGC neuroprotection (100%, OHT-3wk), achieving neuronal density like that observed in sham-operated non-injured control animals. Given that our AAV.CLDN5 conferred EC-specific CLDN5 expression, this robust pro-survival effect is unlikely to result from off-target CLDN5 upregulation in other cell types. This finding highlights the profound harmful effect of a leaky vasculature in glaucoma and the importance of maintaining a tight barrier to control disease pathogenesis. Although the magnitude of vascular leakage in glaucoma may be considered relatively mild, compared to other diseases like age-related macular degeneration, diabetic retinopathy, or ischemic stroke^70^, the chronic nature of glaucoma in which retinal neurons are exposed to a leaky and pro-inflammatory environment over decades is anticipated to be harmful^71, 72^. The strong pro-survival effect of *CLDN5* gene therapy is likely to result from tighter barrier integrity and reduction of extravasation of inflammatory modulators, hence avoiding activation of resident immune cells and glia, which may have damaging effects on neurons^73^. In addition to neuronal survival, AAV.CLDN5 treatment also promoted a significant recovery of visual function as evidenced by marked improvement in optomotor responses. Given that the optokinetic reflex depends on the integrity of RGC axons, these findings indicate that, in addition to RGC soma, CLDN5 gene supplementation successfully protected axons in the optic nerve. Collectively, our findings reveal the importance of DRP1-dependent EC mitochondrial fragmentation on barrier permeability in glaucoma and provide clear evidence that restoring mitochondrial function and CLDN5 homeostasis are valuable strategies to enhance neuroprotection and visual performance.

## MATERIALS AND METHODS

### Experimental animals

All procedures were approved by the animal protection committee of the University of Montreal Hospital Research Centre and followed the Canadian Council on Animal Care guidelines. Experiments included female and male adult Endo-MitoEGFP mice (2-5 months), which were generated by pronuclear injection of the transgenic vector p*Hb9*-MitoEGFP containing the mitochondrial targeting sequence of Cytochrome c oxidase subunit VIII, as previously described^27^, or C57Bl/6 mice (Charles River Laboratories, St-Constant, QC, Canada). Animals were housed in 12h light /12h dark cyclic light conditions, with an average in-cage illumination level of 10 lux, and fed *ad libitum*. Ambient temperature and humidity were maintained at 21-22°C and 45-55%, respectively. All procedures were performed under general anesthesia with isoflurane (2%, 0.8 liter/min in 100% oxygen).

### Mouse glaucoma model

Unilateral elevation of intraocular pressure was induced by a single injection of magnetic microbeads into the mouse eye anterior chamber^28, 29^. Animals were anesthetized and a drop of tropicamide was applied on the cornea to induce pupil dilation (Mydriacyl, Alcon, Mississauga, ON, Canada). A custom-made sharpened microneedle attached to a microsyringe pump (World Precision Instruments, Sarasota, FL) was loaded with a magnetic microbead solution (1.5 µl: 2.4 x 10^6^ beads, diameter: 4.5 µm, Dynabeads M-450 Epoxy, Thermo Fisher Scientific, Waltham, MA). Using a micromanipulator, the tip of the microneedle was gently pushed through the cornea to inject the microbeads into the anterior chamber. The microbeads were immediately attracted to the iridocorneal angle using a hand-held magnet. Sham controls received an injection of PBS. Only one eye was operated on and an antibiotic drop was applied immediately after the surgery (Tobrex, Tobramycin 0.3%, Alcon, Geneva, Switzerland). Intraocular pressure was measured in awake animals before and after the procedure, and biweekly thereafter always at the same time (10 am- 12 pm), using a calibrated TonoLab rebound tonometer (Icare, Vantaa, Finland). For this purpose, a drop of proparacaine hydrochloride (0.5%, Alcon) was applied to the cornea and a minimum of 10 consecutive readings were taken per eye and averaged.

### Retinal immunohistochemistry

Animals were anesthetized and transcardially perfused with ice-cold 4% paraformaldehyde (PFA) in phosphate saline buffer (PBS). Eyes were immediately collected, post-fixed in PFA, and processed to generate cryosections or retinal flat mounts as described^74, 75^. Retinas were incubated at 4°C overnight (cross-sections) or 3 days (whole-mount retinas) in Cytc, (1 μg/μl, BD Biosciences, Franklin Lakes, NJ) or platelet endothelial cell adhesion molecule (CD31, 2 μg/ml, BD Bioscience), followed by fluorophore-conjugated secondary goat anti-mouse Alexa Fluor 594 (1 μg/ml, Life Technologies, Eugene, OR). Sections were rinsed and mounted in antifade solution (SlowFade, Life Technologies) and images were acquired with an Axio Imager M2 optical sectioning microscope (Zeiss, Oberkochen, Germany).

### 3D reconstruction of mitochondrial complexes

Endo-MitoEGFP retinas were incubated in PBS containing isolectin (0.05 µg/µl, (Thermo Fisher Scientific) and TO-PRO-3 was used to label cell nuclei (1 µM, Thermo Fisher Scientific). Images were acquired with a confocal microscope (Leica SP5) using a 100X oil immersion objective (Leica Microsystems Inc., Concord, ON). Images were obtained from each retinal quadrant (dorsal, ventral, nasal, temporal) at the level of the intermediate vascular plexus at 0.5-1.0 mm from the optic nerve head (total area analyzed=5500 µm^2^). Optical sections (z-interval=0.1 µm) were average to reduce noise and the images were captured at a digital size of 1024×1024 pixels (no digital zoom was applied). 3D reconstruction of mitochondria was carried out using the Surface plugin of Imaris software (Bitplane, South Windsor, CT), mitochondrial volume was calculated and normalized relative to vessel volume. The distance outside the nucleus was masked to quantify the volume of mitochondria in the perinuclear area. For vascular density studies, the z stacks containing either the superior, intermediate, or deep plexus were manually separated. Vessel density was quantified using the filament module of Imaris (Bitplane).

### Analysis of mitochondrial superoxide

Endo-MitoEGFP mice received an intravenous injection of the mitochondrial superoxide indicator MitoSOX-Red (33 µM, Life Technologies, Eugene, OR), eyes were collected two hours later, retinas were dissected out and mounted with SlowFade mounting media containing DAPI (4′,6-diamidino-2-phenylindole) (Life Technologies). Images were acquired with an Axio Imager M2 optical sectioning microscope (Zeiss). The outline of EGFP-positive vessels were manually traced and the integrated density (summation of the pixel values in the traced area) was used to measure the MitoSOX-Red-positive fluorescent signal using ImageJ (NIH, Bethesda, MD). Retinal areas with no fluorescence were used as background. The total corrected cellular fluorescence (TCCF) was calculated as follows: TCCF=integrated density – (area of selected cells x mean fluorescence of background readings)^76^.

### Transmission electron microscopy (TEM)

Mice were perfused and fixed using a method designed to reduce morphological damage of metabolically sensitive structures (i.e. mitochondria) in the mouse retina^77^. Briefly, mice were anesthetized and perfused with a fixative solution containing glutaraldehyde (2.5%) and PFA (2%) in sodium cacodylate buffer (0.1 M, pH 7.4) delivered using a controlled syringe pump (20-25 ml, 0.86 ml/min) (Thermo Orion Sage™ Model M361, Thermo Fisher Scientific). Retinas were extracted and post-fixed in the same fixer at 4°C overnight. Samples were then treated with osmium tetroxide (OsO_4_, 1%) for 1 hr at room temperature, dehydrated in increasing concentrations of acetone and embedded in epoxy resin. Ultrathin (120 nm) sections were cut, mounted on 200-mesh copper grids and stained. Pictures were obtained using the AMT XR80C CCD Camera System in a FEI/Philips Tecnai 12 BioTwin 120 kV transmission electron microscopy.

### Vascular permeability assays

i. Fluorescein angiography: For live imaging of vascular leakage, mice were anesthetized and a drop of tropicamide was applied on the cornea to induce pupil dilation (Mydriacyl, Alcon, Mississauga, ON, Canada). Fluorescein (332.32 Da, 5% in 100 µl, Novartis Pharma, Basel, Switzerland) was applied by intravenous injection and images were captured within 5 min of tracer administration using the Micron IV system (Phoenix Research labs, Pleasanton, CA). During the entire imaging session, mice were maintained on a heating pad with a temperature sensor at 37°C. In the visible retinal area, eight regions of interest (ROI) were manually traced (10,000 µm^2^/ROI) and the integrated fluorescence density was measured using Image J to obtain an average fluorescence signal.
ii. Stereological analysis of vascular leakage: Anesthetized mice received a retro-orbital injection of EZ-Link™ Sulfo-NHS-LC-Biotin (0.5 mg/g of body weight, Thermo Fisher Scientific). Eyes were collected 5 min after injection, fixed in ice-cold 4% PFA for 60 min and dissected to generate retinal flat-mounts. Retinas were incubated at 4°C for 3 days in PBS containing Alexa Fluor 488–conjugated isolectin GS-IB4 (Griffonia simplicifolia, 2 μg/ml, Invitrogen, Rockford, IL) and anti-laminin primary antibody (LAMA2, 7 μg/ml, Sigma-Aldrich) followed by an overnight incubation with Alexa Fluor 647–conjugated streptavidin (7 μg/ml, Invitrogen) and Alexa Fluor 488–conjugated donkey anti-rat secondary antibody (2 μg/ml, Invitrogen). Retinas were rinsed and mounted in antifade media (SlowFade, Molecular Probes) and images were obtained with a LSM 900 confocal microscope (Zeiss) using a 20X objective. Z-stack images were acquired using an unbiased stereological approach across the entire retina (175,000 µm^2^/retina) at the level of the superficial, intermediate and deep vascular plexuses. Mean fluorescence intensity of the retinal parenchyma of averaged stack projections was measured with Fiji/ImageJ and normalized to sham-operated retinas.

### Retinal EC preparation

Retinas were isolated and homogenized in physiological buffer (PB:147 mM NaCl, 4 mM KCl, 3 mM CaCl_2_, 1.2 mM MgCl_2_, 5 mM Glucose, 15 mM HEPES) supplemented with protease inhibitors as described^78^. Samples were homogenized using a Dounce homogenizer and centrifuged for 10 min at 3500 x g. The pellet was resuspended in Ficoll solution (20%) in PB, homogenized, and centrifuged 10 min at 25000 x g. The pellet, containing retinal vessels, was resuspended in 15% dextran and overlaid on 20% dextran followed by 15 min of centrifugation at 25000 x g. The resulting pellet of the retinal vessel-enriched preparation was resuspended in PB and passed through a 40 µm cell strainer. Samples were centrifuged 10 min at 25000 x g and final pellets were immediately resuspended in RIPA buffer containing proteases and phosphatase inhibitors for further analysis. All steps were carried at 4°C.

### Western blot analysis

Retinal samples were homogenized with an electric pestle (Kontes, Vineland, NJ) in ice-cold lysis buffer (50 mM Tris pH 7.4, 150 mM NaCl, 1% NP-40, 5 mM Na fluoride, 0.25% Na deoxycholate, and 2 nM NaVO_3_) supplemented with protease and phosphatase inhibitors. Samples were separated in 10% or 12% sodium dodecyl sulfate polyacrylamide gels (SDS-PAGE) and transferred to nitrocellulose membranes (Bio-Rad Life Science, Mississauga, ON). Blots were incubated overnight at 4°C in blocking buffer (10 mM Tris pH 8.0, 150 mM NaCl, 0.1% Tween-20 and 5% bovine serum albumin) containing each of the following primary antibodies: dynamin related protein-1 (DRP1, 1 μg/ml, Cell Signaling Technology, Danver, MA), phosphorylated DRP1 (Ser616) (pDRP1^Ser616^, 1 μg/ml, Cell Signaling Technology), Mitofusin-1 (MFN1, 1 μg/ml, Proteintech, Rosemont, IL), Mitofusin-2 (MFN-2, 1 μg/ml, Cell Signaling Technology), Optic protein atrophy-1 (OPA1, 1 μg/ml, Cell Signaling Technology), Claudin-5 (CLDN5, 1 μg/ml, ThermoFisher Scientific), occludin (0.25 μg/ml, Invitrogen), vascular endothelial-cadherin (VE-Cadh, 0.2 μg/ml, R&D Systems, Minneapolis, MN), phosphorylated VE-Cadherin (Tyr731) (pCadh^Tyr731^, 1 μg/ml, OriGene Technologies, Rockville, MD), phosphorylated VE-Cadherin (Tyr658) (pCadh^Tyr658^, 1 μg/ml, Millipore, Temecula, CA), or β-actin (0.5 μg/ml, SIGMA-Aldrich, St Louis, MO). Membranes were washed and incubated in peroxidase-linked anti-mouse, anti-rabbit or anti-goat secondary antibodies (0.5 μg/ml, GE Healthcare, Mississauga, ON). Blots were developed using a chemiluminescence reagent followed by analysis using the ChemiDoc MP System (Bio-Rad). Densitometric analysis was performed using Image Lab capture and analysis software from a series of three independent western blots each carried out using retinal samples from distinct experimental or control groups.

### Pharmacological DRP1 inhibition

For pharmacological DRP1 inhibition, mice received daily intraperitoneal injections of the mitochondrial division inhibitor-1 (Mdivi-1, 3-[2,4-dichloro-5-methoxyphenyl]-2-sulfanyl-4[3H]-quinazolinone, 20 mg/kg, Enzo Life Sciences, Farmingdale, NY).

### EC-targeted gene delivery

The following AAV serotype 9 vectors were administered by tail vein injection for selective gene expression in ECs three weeks prior to glaucoma induction (100 µl total volume): i) AAV.DRP1^K38A^ (2.0×10^13^ GC/ml), encoding dominant negative DRP1 (K38A mutation)^43^ tagged with HA, ii) AAV.CLDN5 (3.7×10^13^ GC/ml), encoding CLDN5 tagged with mCherry, or iii) AAV.Ctl (5.4×10^12^ GC/ml) encoding only mCherry (Vector Biolabs Malvern, PA). All expression cassettes were driven by the EC-specific Ple32 (*CLDN5*) minipromoter^44^.

### Quantification of RGC survival

Mice were transcardially perfused with 4% PFA, retinas were dissected out and incubated for three days with an antibody against RBPMS (0.84 µg/ml, PhosphoSolutions) followed by an Alexa Fluor 647-conjugated secondary antibody (4 μg/ml, Invitrogen). Retinas were washed and mounted using an antifade reagent (SlowFade, Life Technologies)^79, 80^. Images were obtained using an Axio Imager M2 optical sectioning microscope (20X objective, Zeiss) equipped with an automated stage for X-, Y-, and Z-axis movement, a color camera (Axiocam 509 mono, Zeiss), and image analysis software (Zen, Zeiss). We used an unbiased stereological approach based on systematic uniform random sampling from 3D-dissectors (stacks) across the entire retina, and images were acquired using identical exposure time and gain settings for all experimental and control groups as described^24, 59^. RGC soma were quantified using a custom-made quadrant dissector in Fiji/ImageJ.

### Optomotor response assay

Visual acuity was evaluated by measuring the optomotor reflex using the OptoMotry virtual reality system (Cerebral Mechanics Inc., Medicine Hat, AB, Canada), which allows the quantification of visuomotor behaviors in response to visual stimuli in glaucoma^40, 59^. The animals were placed on an elevated platform in the center of a testing arena with walls composed of computer monitors displaying a rotating vertical black and white sinusoidal grating pattern. The staircase method was used to determine the spatial frequency applying sinusoidal steps. Both contrast (100%) and rotation speed (12°/s) were kept constant. Cameras were used to record the animals, and an observer blinded to treatment monitored their tracking behavior. A behavioral response was considered positive when the motor response (head movement) was concordant with the direction of the visual stimulus (moving bars). Individual scores from each mouse were collected before and after each treatment.

### Statistical analyses

Data analysis was always carried out blinded by third party concealment of treatment using uniquely coded samples. Statistical analysis was performed with Prism 10 version 10.4.1 (GraphPad, San Diego, CA). We evaluated all cohorts with normality (Shapiro-Wilk test) and variance (F-test) tests. Values were compared by means of two-tailed Student’s t-test or Mann-Whitney U test, where appropriate. For multiple comparisons, we used Analysis of Variance (ANOVA) followed by Dunnett’s or Tukey’s test, where appropriate. A p value ≤0.05 was considered significant. The number of animals used in each experiment are indicated in the figure legends. All values are provided as the mean ± standard error of the mean (S.E.M.), and individual values are presented in each graph.

## ACKNOWLEDGEMENTS

The authors thank Drs. Yoko Ito and Yukihiro Shiga for helpful comments, advice, and assistance during this project; Aurélie Cleret-Buhot (Imaging platform) at the Université de Montréal Hospital Research Center for technical support; and Jeannie Mui and Weawkamol Leelapornpisit (Electron Microscopy Facility, McGill University) for assistance with sample preparation, microscope operation, and data collection.

## AUTHOR CONTRIBUTIONS

Conceptualization: JLCV, NB, IAVP, HQ, ADP. Methodology: JLCV, NB, IAVP, FD, CVV, HQ, ADP. Investigation and visualization: JLCV, NB, IAVP, FD, HQ, ADP. Writing and editing: JLCV, NB, IAVP, CVV, HQ, ADP. Supervision: ADP. Funding acquisition: ADP. All authors reviewed and approved the manuscript.

## AVAILABILITY OF DATA AND MATERIALS

All data generated or analyzed during this study are included in this published article. Materials are available upon request.

---

**Supp. Figure 1.**
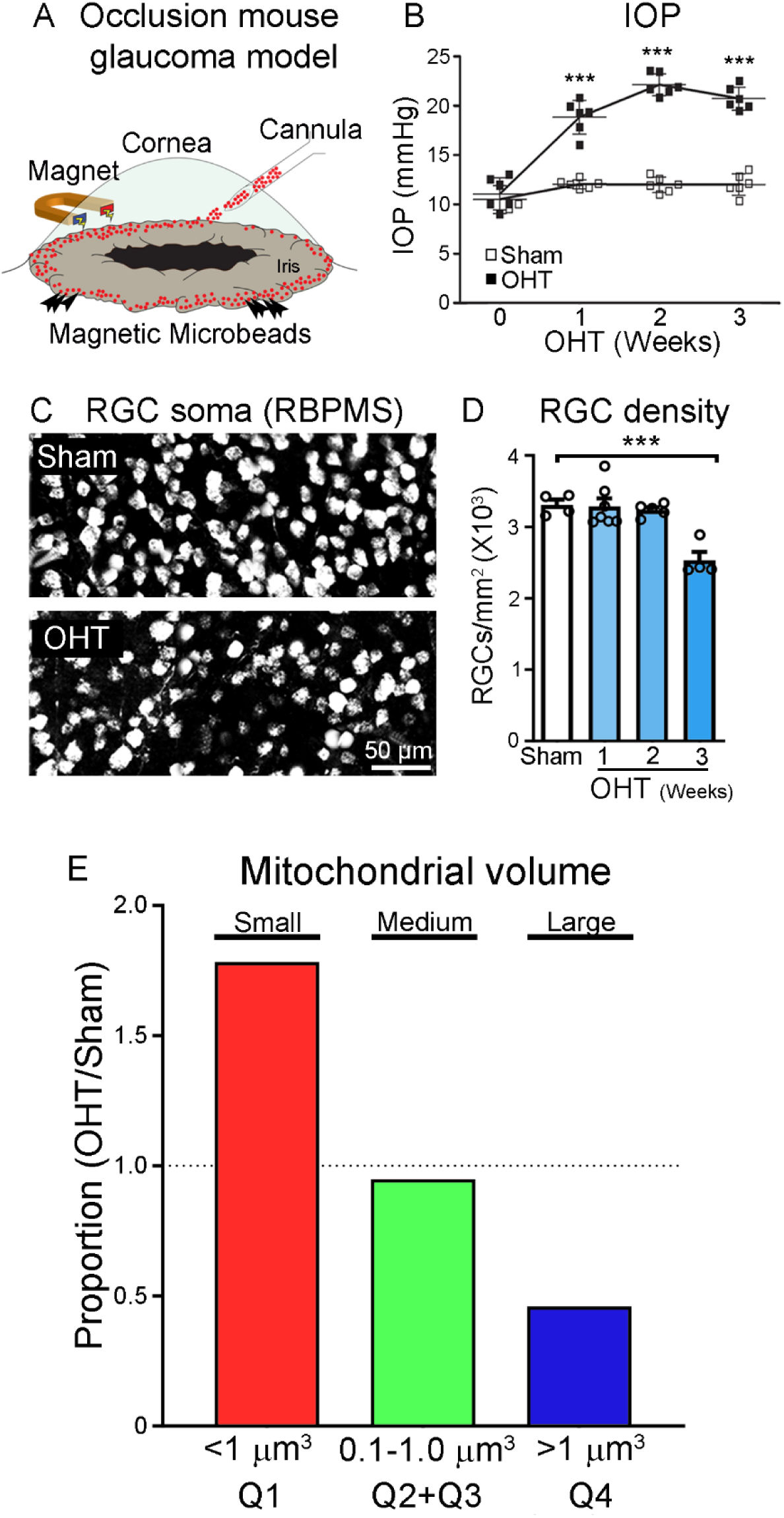
Intracameral injection of magnetic microbeads increases intraocular pressure and causes progressive RGC loss. (A, B) Schematic of the mouse glaucoma model induced by intracameral injection of magnetic microbeads results in a gradual increase of intraocular pressure (IOP) causing ocular hypertension (OHT) (N=8 mice/group, two-way ANOVA with Sidak’s multiple comparison post hoc test, ***p<0.001). (C) Representative images of RBPMS-labeled RGCs in sham and glaucomatous mice at three weeks after magnetic microbead injection. (D) Elevated IOP leads to significant RGC loss starting at three weeks after glaucoma induction (N=4-8 mice/group, ANOVA, Tukey’s Multiple Comparison *post-hoc* test, ***p<0.001). (E) EC mitochondrial volume distribution in each quartile in OHT eyes normalized to sham eyes: i) Q1: lower quartile defined as small (<0.1 µm^3^), ii) Q2+Q3: inter-quartiles defined as medium (0.1-1 µm^3^), and iii) Q4: upper quartile defined as large (>1 µm^3^). The cartoon in this figure were generated with BioRender (https://biorender.com).

**Supp. Figure 2.**
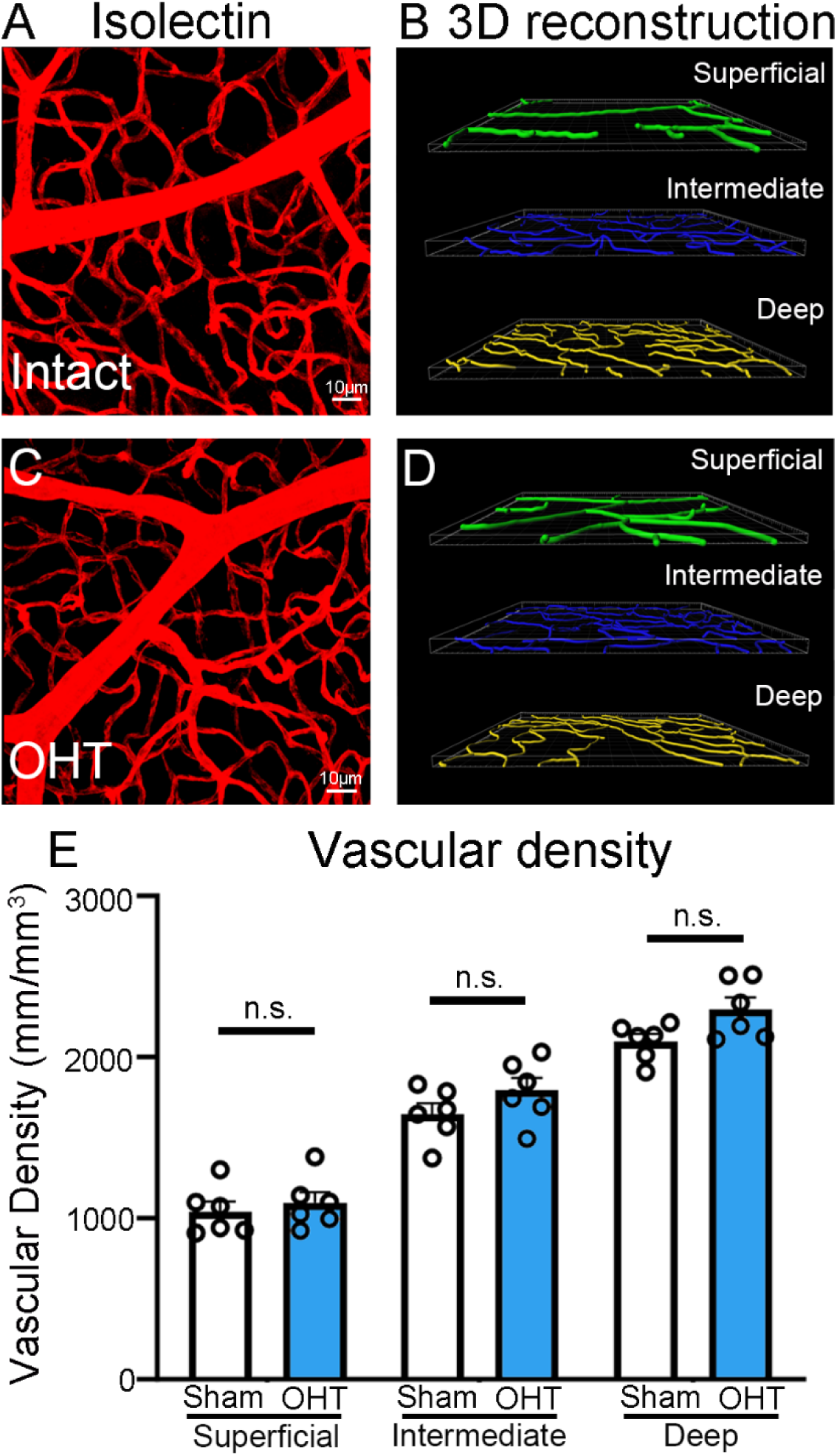
Increased intraocular pressure does not reduce capillary density. (A-D) Whole-mounted retinas showing vessels labeled with isolectin (A, C) and 3-dimensional reconstruction of the capillary network in the superficial, intermediate, and deeper plexuses (B, D) show similar vascular densities in intact versus glaucomatous eyes (OHT-3wks). (E) Quantification of the retinal capillary density in the superficial, intermediate, and deep plexuses confirms that there is no vessel loss after OHT induction (N=6 mice/group, ANOVA, Tukey’s multiple comparison *post-hoc* test, n.s.: not significant).

**Supp. Figure 3.**
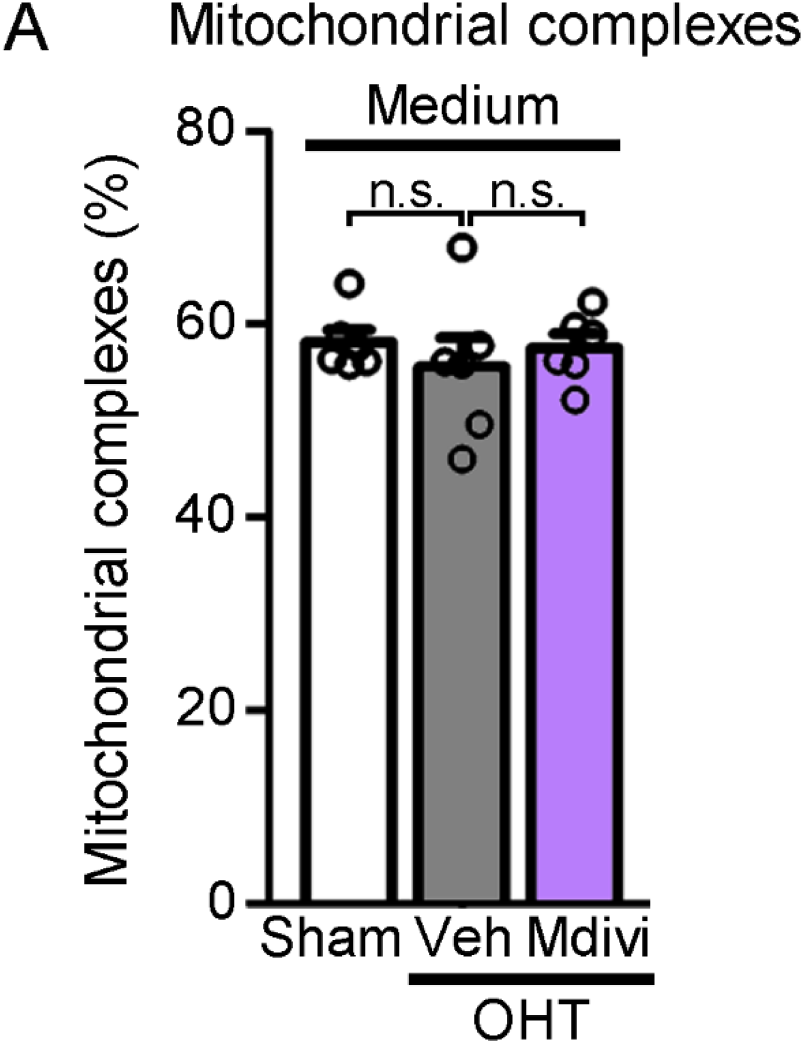
The number of medium-sized mitochondrial complexes does not change with OHT. (A) Quantitative analysis of 3D-reconstructed EC mitochondrial complexes show that there are no statistically significant differences in the number of medium size mitochondrial complexes between sham, OHT + vehicle, OHT + Mdivi-1-treated retinas (N=6 mice/group, ANOVA, Tukey’s multiple comparison *post hoc* test, n.s.: not significant).

**Supp. Figure 4.**
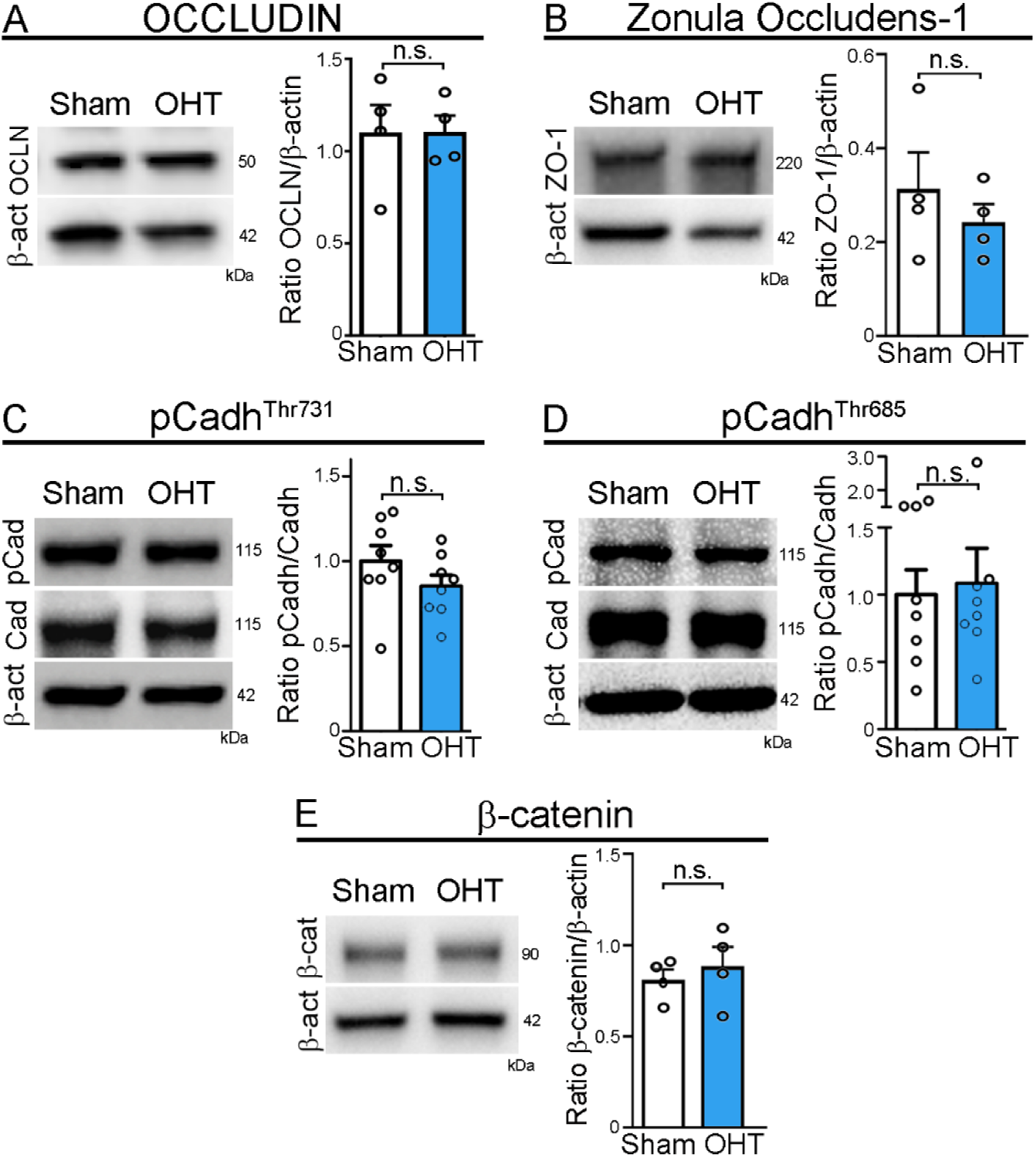
Levels of other tight and adherens junctions remain unchanged during OHT. Western blot analysis of fresh retinas revealed that OHT does not lead to statistically significant changes in the tight or adherens junctions: (A) occludin (OCLN), (B) ZO-1, (C) phosphorylated VE-Cadherin at Threonine 731 (pCadh^Thr731^), (D) phosphorylated VE-Cadherin at Threonine 685 (pCadh^Thr685^), or (E) b-catenin (N=5 mice/group, two-tailed Student’s t-test, n.s.: not significant).

**Supp. Figure 5.**
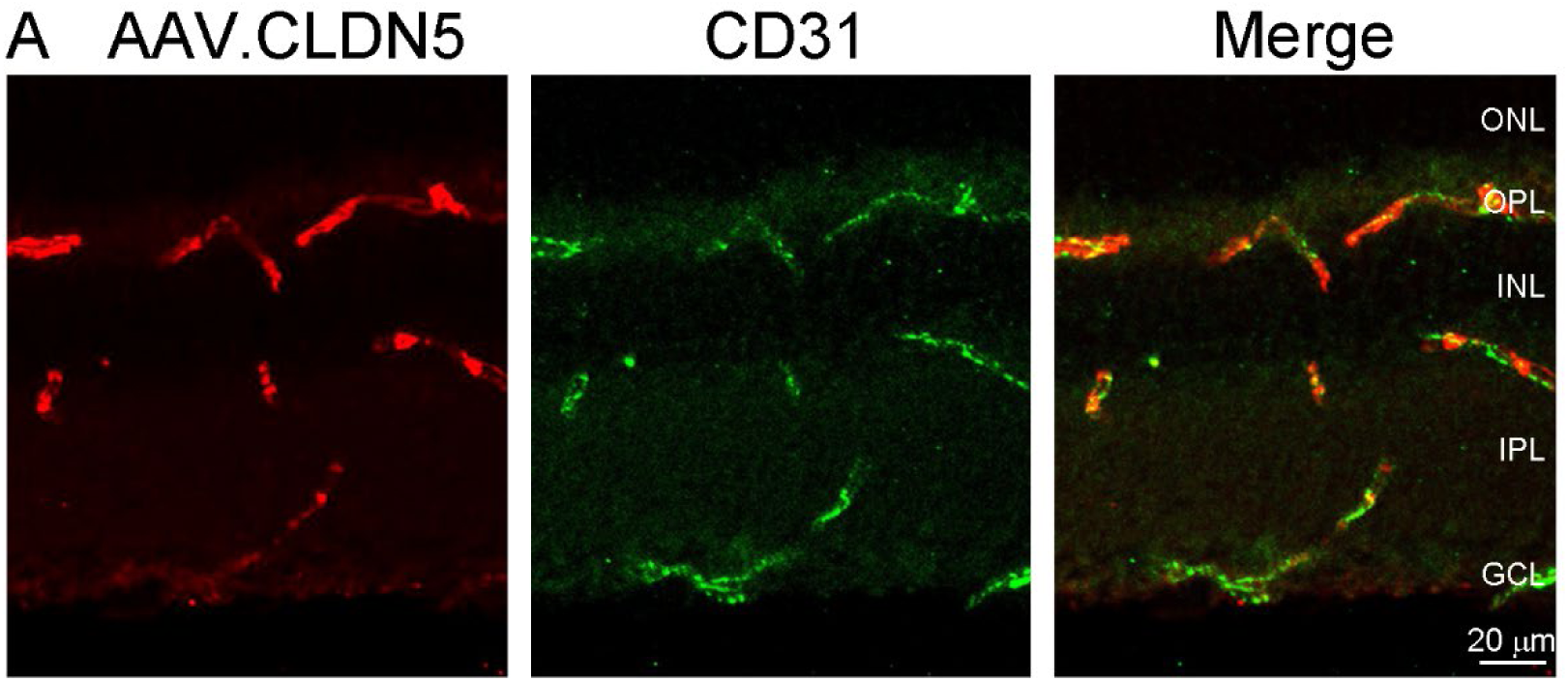
EC-specific AAV-mediated CLDN5 expression. (A) Representative retinal cross-section showing EC-specific expression of AAV-mediated CDLN5 (red) after four weeks of tail vein injection of AAV.CLDN5. ECs were visualized with the EC-specific marker CD31 (green). No other cell types were labeled after tail vein injection of AAV.CLDN5.

